# Potential role of microRNAs in regulating transcriptional profile, and sculpting development and metabolism in cavefish

**DOI:** 10.1101/2024.01.30.578051

**Authors:** Tathagata Biswas, Huzaifa Hassan, Nicolas Rohner

## Abstract

*Astyanax mexicanus*, a species with both surface-dwelling and multiple cave-dwelling populations, offers a unique opportunity to study repeated adaptation to dark and resource-scarce environments. While previous work has identified large-scale changes in gene expression between morphs even under identical laboratory conditions, the regulatory basis of these expression differences remains largely unexplored. In this study, we focus on microRNAs (miRNAs) as key regulators of gene expression to understand cavefish adaptation nuances. Our analysis identified 683 miRNAs, which not only surpasses the number documented in related species but also provides the first comprehensive catalog of miRNAs for this species. We identified a unique subset of differentially expressed miRNAs common to all studied cave-dwelling populations, potentially orchestrating the nuanced gene expression patterns required for survival in the challenging cave milieu. Gene Ontology analysis of the predicted miRNA targets revealed involvement in developmental and metabolic pathways that are pivotal for thriving in nutrient-limited environments, such as the regulation of neuromast migration. Moreover, our study provides evidence for miRNA influence on circadian rhythm and oxidative stress response, both essential adaptations for the cave-dwelling lifestyle. The comprehensive miRNA catalog generated will guide future investigations into the intricate world of miRNA-mediated evolution of complex traits.

## Introduction

The fish species *Astyanax mexicanus* is an exceptional animal model for understanding adaptation to dark and nutrient-limited environments. This species exists as river-dwelling surface fish and cave-dwelling populations that remain fertile with each other and can be kept in the laboratory. Less than 200,000 years ago, the ancestral surface fish colonized multiple caves in the Sierra de El Abra and neighboring mountain regions of Mexico, through multiple independent events.^1, 2^ Inside each of these caves, under similar environmental pressures, *Astyanax* evolved a range of similar traits that enabled the cave morphs to not only survive but thrive in the dark and therefore biodiversity and nutrient-limited cave environment.^3-9^ As a consequence, the cave morphs show significant and heritable changes to their transcriptomic landscape, when compared to the extant surface fish.^5, 10-13^ While the role of cis-regulatory enhancers in these transcription changes has been studied,^13^ the exploration of other mechanisms of transcriptional regulation remains less understood. One particularly important mechanism that warrants further investigation is the role of non-coding RNAs in modulating gene expression in *Astyanax mexicanus*. A class of non-coding RNA that plays an important role in shaping gene expression profiles are microRNAs (miRNAs). The miRNAs are a class of small endogenous non-coding RNA molecules, typically averaging around 18-25 nucleotides in length, which act post-transcriptionally to regulate gene expression.^14-17^ Exported into the cytoplasm from the nucleus as short hairpin structures, one of the strands of a mature miRNA is eventually loaded into the RNA-induced silencing complex. The silencing complex most frequently targets the 3’UTR region of mRNAs through complementary base pairing and fine-tunes gene expression through translational repression or mRNA degradation.^18-20^ A single miRNA can target multiple mRNAs, and conversely, a single mRNA can be regulated by many different miRNAs, amplifying the regulatory potential of miRNA-mediated regulation of gene expression.^21^ However, despite being extensively studied in multiple model organisms, miRNA’s potential role in altering the transcriptomic profile of cavefish remains unexplored. ^22-25^ In this study, we focused on 24-hour-post-fertilization (hpf) embryos to isolate and annotate miRNAs expressed in four different morphs of *Astyanax mexicanus*: surface fish, and three independently or parallelly evolved populations of cavefish (Pachón, Tinaja, and Molino). Using miRDeep2 package^26^ to call for novel miRNAs in *Astyanax mexicanus*, we identified as many as 683 different miRNAs, surpassing comparable studies in zebrafish.^27-29^ Our comparative analysis of miRNA expression between three cave-dwelling and surface-dwelling populations of *Astyanax mexicanus* identified a cave-specific set of differentially expressed miRNAs and their putative 3’UTR targets. Gene Ontology analysis of these targets suggests that miRNA regulation may play a critical role in cave adaptation by influencing developmental and metabolic pathways. This study not only enriches the miRNA catalog for this species but also underscores the potential of miRNA-mediated regulation in the evolutionary response to extreme environments.

## Materials and Methods

### Small RNA isolation and sequencing

We collected 50 embryos of each morph of *Astyanax mexicanus*, at the 24hpf developmental stage. After a 1xPBS wash, samples were flash-frozen and homogenized. The small RNA population from each morph was isolated using the mirVana miRNA Isolation Kit (Catalog number: AM1560) following manufacturer’s instructions.

Small RNA-seq libraries were generated from 50ng of enriched small RNA, as assessed using the Bioanalyzer (Agilent). Libraries were made according to the manufacturer’s directions for the TruSeq Small RNA Library Prep Kit (Illumina, Cat. No. RS-200-0012). Resulting libraries were size selected for 145-160bp fragments using Novex 6% TBE gels (Invitrogen, Cat. No. EC6265BOX) to retain only the small RNA inserts. Size selected libraries were checked for quality and quantity using the Bioanalyzer (Agilent) and Qubit Fluorometer (Life Technologies). Libraries were pooled, re-quantified and sequenced as 75bp single read on an Illumina NextSeq 500 high-output flow cell utilizing RTA and current instrument software versions at the time of processing.

### Adapter removal and filtering of reads

The ‘TGGAATTCTCGGGTGCCAAGG’ adapter was removed for all samples of small RNA-seq using Cutadapt^30^ (v2.5). Reads without an adapter and shorter than 18bp were discarded. We excluded reads aligning to rRNAs, scaRNAs, snoRNAs, and snRNAs. Additionally, reads aligning to tRNAs were removed. As annotated tRNAs for *Astyanax mexicanus* 2.0 are unavailable, we used tRNAscan-SE^31^ (v2.0.6) to search for *Astyanax mexicanus* 2.0 tRNAs using Danio rerio (GRCz11) tRNAs from GtRNAdb (the genomic tRNA database).

### Novel miRNA Identification

The reads remaining after filtering (rRNAs, scaRNAs, snoRNAs, snRNAs and tRNAs) from all samples were concatenated and the miRDeep2 ^26^ (v3) tool was used to identify novel miRNAs. First, the mapper.pl module was used to align the reads to *Astyanax mexicanus* 2.0 genome from NCBI (Assembly GCF_000372685.2). Next miRDeep2.pl module was run on the mapped arf file using the reference genome and known mature *Danio rerio* miRNAs from miRBase (v22) as supporting miRNAs from related species. The resulting novel miRNAs were further filtered based on the following criteria: mirDeep2 score >= 10, mature read count >=10, and a significant randfold p-value. Novel miRNAs with a matching seed sequence to *Danio rerio* miRNAs from miRBase were retained, even if they did not meet the above criteria. CD-HIT ^32^ (v4.6) tool was used to collapse the redundant novel miRNAs with 100% identity.

### miRNA conservation analysis

To assess the conservation of novel miRNAs across known miRNAs from all species in miRBase, the mature sequence of each novel miRNA was input into BLASTn (v2.10.1+) to find similar sequences against all the know miRNAs. The default E-value threshold of 10 was used to report all hits.

### Quantification of miRNAs and Differential Expression Analysis

To quantify the expression of each novel miRNA in all the samples we used quantifier.pl module in miRDeep2 and aligned the sequenced reads to the novel precursor miRNAs using the default parameters.

Differentially expressed miRNAs were determined using R package edgeR^33^ (v3.22.3). The resulting p-values from the differential expression analysis were adjusted using the Benjamini-Hochberg method with the R function p.adjust. miRNAs with an adjusted p-value less than 0.05 and a fold change of at least 2 were considered as differentially expressed.

### miRNA-mRNA target identification

miRanda^34^ (v3.3a) was used to identify the miRNA-mRNA targets using a score threshold of > 80 for scores and energy threshold < -14 kcal/mol. The novel miRNAs identified were checked for targets against the 3’ UTRs of 11,318 *Astyanax mexicanus* 2.0 genes.

### mRNAseq dataset processing

Pachón and surface fish mRNA-seq datasets for two time points: 24hpf and 36hpf were downloaded from SRA database: SRP045680. The reads were aligned to *Astyanax mexicanus* 2.0 reference genome, NCBI assembly GCF_000372685.2 using STAR^35^ (v2.7.3a). The gene model retrieved from Ensembl, release 102 was used to generate gene read counts. The transcript abundance ‘TPM’ (Transcript per Million) was quantified using RSEM^36^ (v1.3). Differentially expressed genes were determined using R package edgeR^33^ (v3.38.4). Prior to differential expression analysis, low-expression genes were filtered out based on a cutoff of 0.5 CPM (Counts Per Million) in at least one library. The resulting p-values were adjusted with Benjamini-Hochberg method using R function p.adjust. Genes with an adjusted p-value < 0.05 and a fold change of 2 were considered as differentially expressed.

### Gene Ontology (GO) analysis

Gene functional enrichment analysis or GO analysis was performed using a custom script built over R package clusterProfiler^37^ (v4.4.4). *Astyanax mexicanus* Gene–GO terms retrieved from Ensembl BioMart were used to identify overrepresented GO terms in the differentially expressed genes compared to the background list of all genes.

## Results

### Identification and annotation of miRNAs in *Astyanax mexicanus*

To establish a comprehensive repository of miRNAs in *A. mexicanus*, we sequenced, identified, and annotated small miRNA populations from surface fish and three distinct cavefish populations – Pachón, Tinaja, and Molino. Concurrently, we aimed to investigate the potential impact of miRNAs on the development of cave traits. One of the most obvious developmental differences between cave and surface fish occurs around 24hpf, when the eye in cavefish starts to regress.^9, 38, 39^ Thus, we focused on 24hpf embryos to explore differentially expressed miRNAs between cavefish and surface fish.

To enhance the efficiency of miRNA identification, we specifically isolated and sequenced only small RNAs (<200nt) instead of total RNA from the embryos (Figure 1a). After adapter trimming, the small RNA population between read lengths 15bp - 40 bp was found to mostly enrich for reads in the range of 21-30bps (Figure 1b). This was in line with the known range of miRNA lengths in zebrafish (Figure 1c). Subsequently, the reads were aligned to the *Astyanax mexicanus* 2.0 genome and the trimmed files of all the samples were merged to call for novel miRNAs using miRDeep2^26^ with default parameters and zebrafish miRNA from miRBase as the supporting miRNA from related species. After removing redundancies using established pipelines, we identified and annotated a refined list of 683 miRNAs (Supplementary Table 1) from an initial collection of 1,452. Of these 683 miRNAs, 681 were conserved across different organisms and had a hit in the miRBase (Supplementary Figure S1). Of note, we identified more miRNAs in cavefish than the 374 miRNAs currently cataloged for zebrafish in the miRBase database.

**Figure 1:**
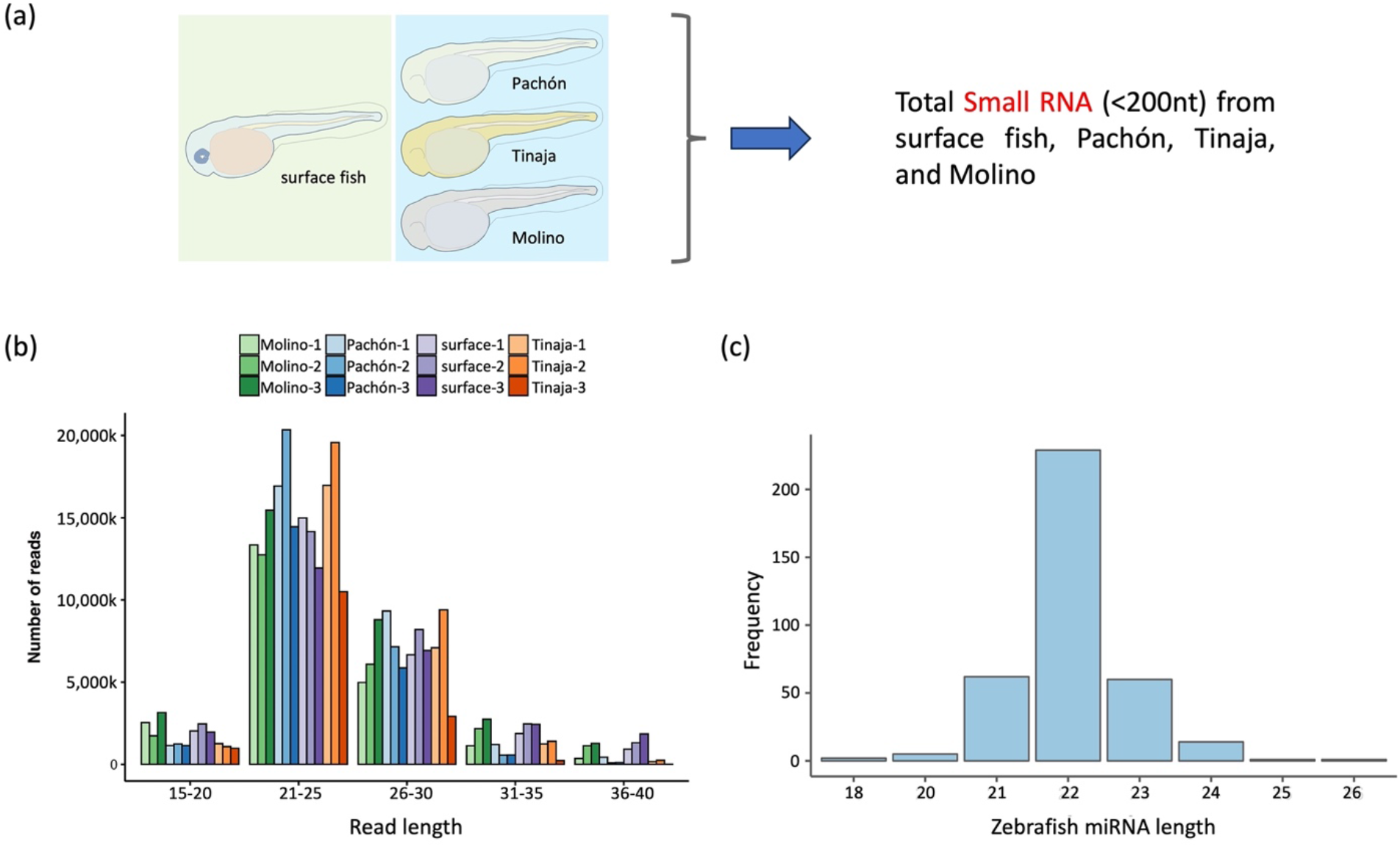
Identification of miRNAs in *Astyanax mexicanus*. (a) Graphical representation of the four morphs – surface fish, Pachón, Tinaja, and Molino at 24hpf, from which the small RNA (<200nt) population was collected. (b) Distribution of small RNA reads across read length of 15 – 40bp. (c) Size distribution of all the zebrafish miRNAs available on miRBase.

### Differential, and cave-specific, expression of miRNAs across four morphs

Small non-coding miRNAs are recognized for their role in facilitating evolutionary adaptation in organisms to environmental stresses.^40-42^ To explore any such correlation in cavefish development and adaptation, we investigated miRNAs which were differentially expressed in the cave morphs when compared to the surface fish. Comparing the miRNA expression profile of each of the cave morphs (Pachón, Tinaja, and Molino) with that of the surface fish, at 24hpf, revealed both up- and downregulated miRNAs (Figure 2a). Pachón and Tinaja had similar numbers of differentially expressed miRNAs, totaling 203 and 196 respectively, while Molino exhibited the highest abundance of differentially expressed miRNAs among the three cave morphs, totaling 238. The highest number of upregulated miRNAs was observed in Molino, followed by Pachón and Tinaja, in comparison to surface fish. Conversely, the highest number of downregulated miRNAs was observed in Tinaja, followed by Pachón and Molino.

**Figure 2:**
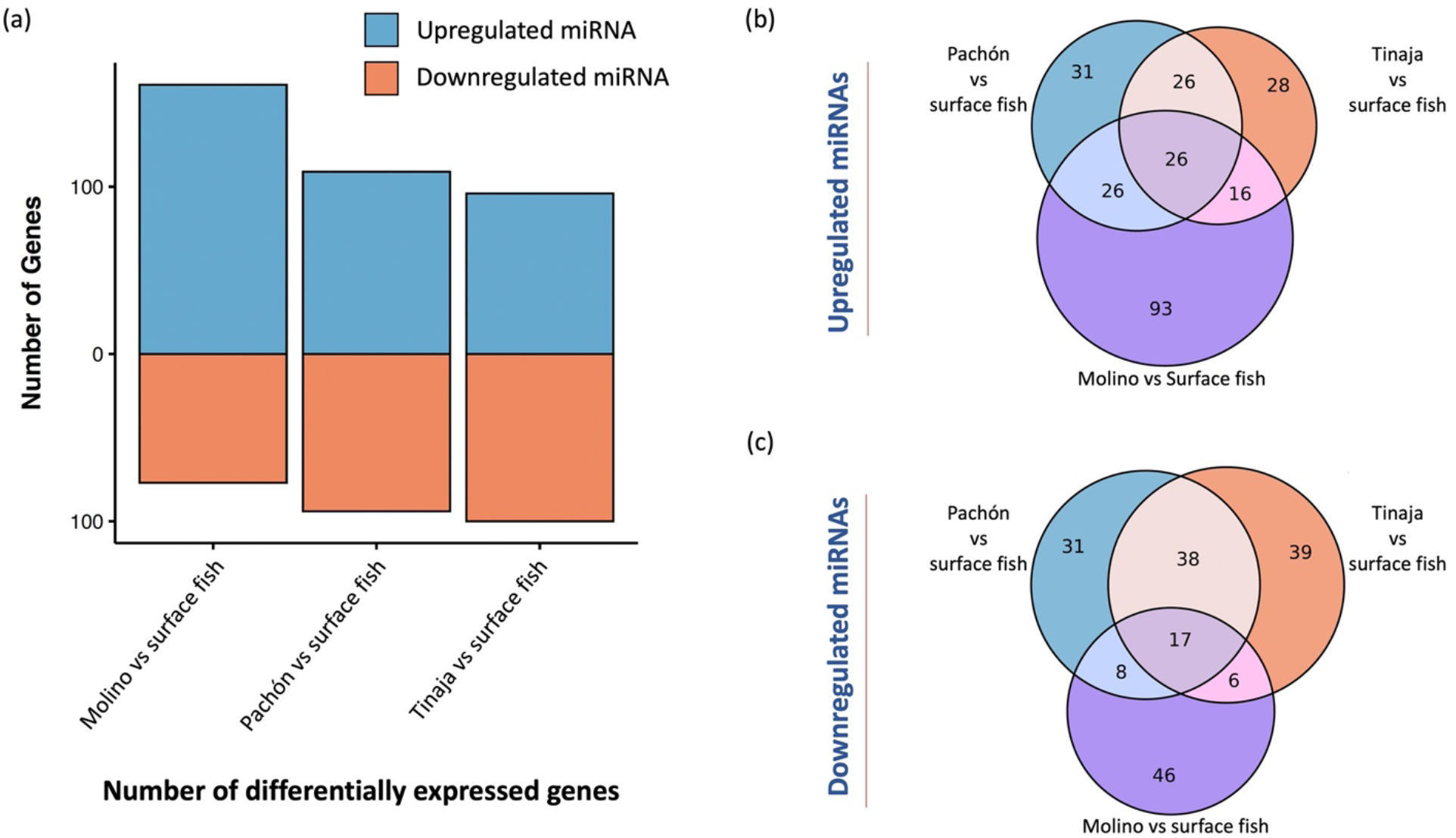
Differentially expressed miRNAs in cave morphs when compared to surface fish. (a) Abundance of differentially expressed miRNAs across Molino, Pachón, and Tinaja (arranged in order of abundance). (b) Venn diagram depicting extent of overlaps in miRNAs upregulated in each of the cave morphs, compared to surface fish. The central overlap of 26 miRNAs depicts the cave-specific miRNAs, upregulated in all the three cave morphs, with respect to surface fish. (c) A similar Venn diagram as ‘(b)’, depicting extent of overlaps in miRNAs downregulated in each of the cave morphs, compared to surface fish.

Next, we explored not just morph-specific, but also cave-specific differentially expressed miRNAs. We defined cave-specific miRNAs as those miRNAs that are up- or downregulated across all the three cave morphs when compared to surface fish. We argued that cave-specific miRNAs would be strong candidates for underlying early adaptation in cave morphs. From the list of all the differentially expressed miRNAs, 26 miRNAs were upregulated (Figure 2b) and 17 were downregulated (Figure 2c) across Pachón, Tinaja, and Molino, when compared to surface fish. The set of 26 and 17 miRNAs represents the first identified cave-specific, non-coding miRNAs differentially expressed across three different cave morphs, with respect to surface fish.

### Putative 3’UTR targets of miRNA in *Astyanax mexicanus*

To further understand the functional implications of these cave-specific miRNAs, we turned our attention to their putative targets. miRNAs mostly target the 3’UTR region of genes. Therefore, to predict putative miRNA target genes in *Astyanax mexicanus*, we used the miRanda pipeline. The miRanda pipeline utilizes annotated 3’UTR of genes to score and predict target mRNAs through dynamic-programming alignment and thermodynamics for miRNAs. It is to be noted that in *A. mexicanus*, only less than half of the annotated genes have an annotated 3’UTR, specifically, 11,318 genes out of a total of 27,420 genes with the annotated 3’UTRs spanning over a considerable range of lengths, with a mean of around 2-kb (Supplementary Figure S2). Therefore, restricting our predictions within the list of annotated 3’UTRs, we identified the putative target mRNAs for each of the 683 miRNAs generated the target list with miRanda scores (see Methods section). As expected, each of the miRNAs had multiple 3’UTRs as putative targets, thereby also making each gene a target of multiple miRNAs (Supplementary Figure S3, Supplementary Table 2).

### Cave-specific miRNAs and Gene Ontology analysis of their targets

Next, we used the list of predicted targets to identify potential pathways through which miRNAs could regulate cave adaptation. As mentioned earlier, the miRNAs that show differential expression across all three cave morphs, when compared to surface fish, are likely to repeatedly contribute to the development of cave-specific traits and as such have a higher likelihood of playing important roles in the development of cave features (cave-specific miRNAs). Therefore, we focused on putative targets of these cave-specific miRNAs, and for each of the 26 upregulated and 17 downregulated miRNAs (Figure 2b,c), from our database of predicted targets, we shortlisted the top 10 target genes based on corresponding miRanda scores and processed them through GO term analysis. For the set of 26 cave-specific upregulated miRNAs and each of their top 10 target genes (Supplementary Table 3), the putative biological pathways affected were “negative regulation of vasculature development”, “regulation of Rho protein signal transduction”, “neuromast primordium migration”, and “ionotropic glutamate receptor signaling pathway” (Figure 3a). The presence of a neuromast-associated pathway in this list, suggesting early alteration of neuromast development, is in line with what is known about cavefish evolution. To compensate for their loss of visual capacity inside the caves, the cavefish exhibit more numerous, larger, and more sensitive neuromast population, as compared to surface fish.^43, 44^ Similarly, the presence of glutamate receptor signaling pathway^45, 46^ among the GO terms potentially suggests an early window into understanding differential neurotransmission in cave morphs compared to surface fish.

**Figure 3:**
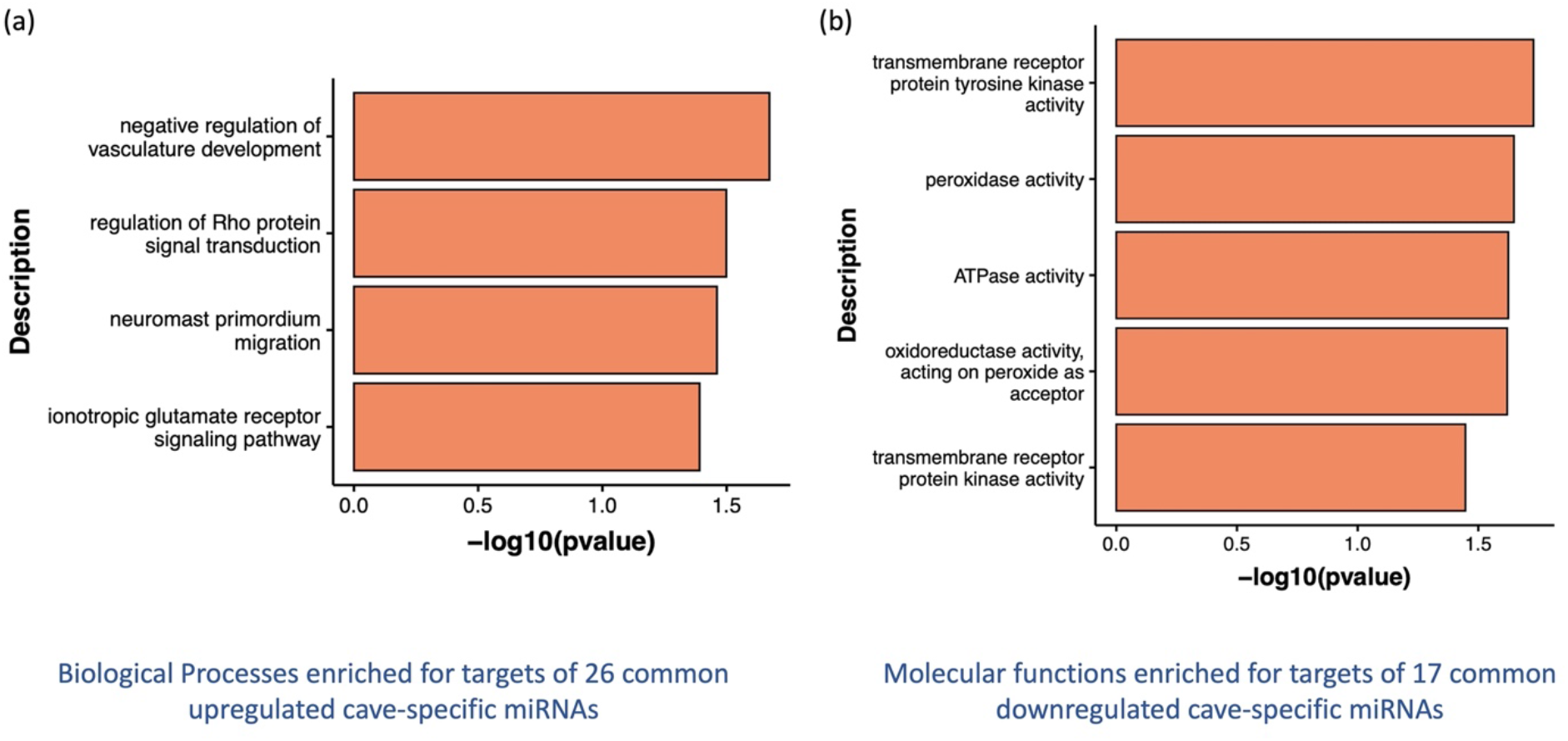
Pathways enriched for top 10 target genes of cave-specific miRNAs. (a) Biological processes enriched for the targets of 26 cave-specific upregulated miRNAs. (b) Molecular functions of target genes of the 17 cave-specific downregulated miRNAs.

Next, we analyzed the top 10 targets for each of the seventeen miRNAs downregulated in cavefish (Supplementary Table 4), conducting a comparable Gene Ontology (GO) term analysis. Most of the biological processes enriched for cell cycle regulation were not directly associated with adaptation to cave or extreme environments (Supplementary Table 5). However, the molecular functions of the listed targets, which included genes like *prdx5* and *prdx4*, enriched pathways associated with oxidative metabolism like “peroxidase activity” and “oxidoreductase activity, acting on peroxide as acceptor” (Figure 3b). It is known that long periods of starvation and hyperglycemia, both contribute to increased oxidative stress in organisms.^47, 48^ Cavefish are adapted to be resilient to long periods of starvation and are also hyperglycemic,^3, 7, 49^ thereby potentially necessitating enhanced antioxidant capacity in cavefish to protect against oxidative stress.^50^

### Cross-validation targets with mRNAseq dataset

While silencing gene expression, apart from repression of translation, miRNAs also work through mRNA degradation. To that effect, we investigated if the predicted targets of cave-specific miRNAs were differentially expressed at comparable embryonic stages. Here, instead of using top 10 targets, we utilized all the genes which were predicted to be targeted by cave-specific miRNAs. As a sample set for analysis, we utilized an available mRNAseq dataset of Pachón – as representative for cavefish, and surface fish – as control.^25^ From this dataset, we merged differentially expressed genes from Pachón versus surface fish at two timepoints, 24hpf and 36hpf. Our approach considered the time-dependent nature of miRNA action, wherein it takes a few hours for the miRNA to be expressed at a certain time point to exhibit its effect.^51, 52^ Merging the two data points, we found 4,088 unique differentially expressed genes in Pachón, when compared to surface fish. Out of these differentially expressed genes, a total of 1,598 were from the list of predicted targets of the cave-specific miRNAs (Figure 4a), amounting to 39.1% of the differentially expressed genes. Subsequently, we employed GO term analysis over these 1,598 genes to explore the associated biological pathways. Notably, several of the essential pathways commonly associated with cave adaptation were present in the list (Figure 4b, Supplementary Table 6). For example, we observed enrichment of pathways associated with “regulation of circadian rhythm”. Genes like *per2, hcrtr2*, and *nptx2b*, which were already listed as targets of the cave-specific miRNA set, were also differentially expressed (Figure 4c), contributing to the enrichment of the circadian pathways. While both *per2* and *hcrtr2* are known to be important for cave adaptation through the regulation of circadian rhythm, and sleep cycle,^4, 53^ *nptx2b* is known to function as a modulator of circadian synaptic variations.^54^ Additionally, *per2* is also known to regulate lipid metabolism through regulation of peroxisome proliferator-activated receptor γ.^55, 56^ We also observed an enrichment of neural pathways and pathways associated with immune systems. While cavefish neural projections and activity differs considerable from that of surface fish, owing to their modified sensory inputs system, the immune system of cavefish is also different from that of surface fish, where cavefish exhibit decreased innate immune response compared to surface fish.^6^ Interestingly, even in an *in vitro* set-up, immune response has been observed to be one of the differentially regulated pathways between surface fish and cavefish cells.^57^ Modifications in both of these overarching pathways plays a major role in enabling the cave morphs to thrive in caves highlighting the potential role of miRNAs in shaping these traits.

**Figure 4:**
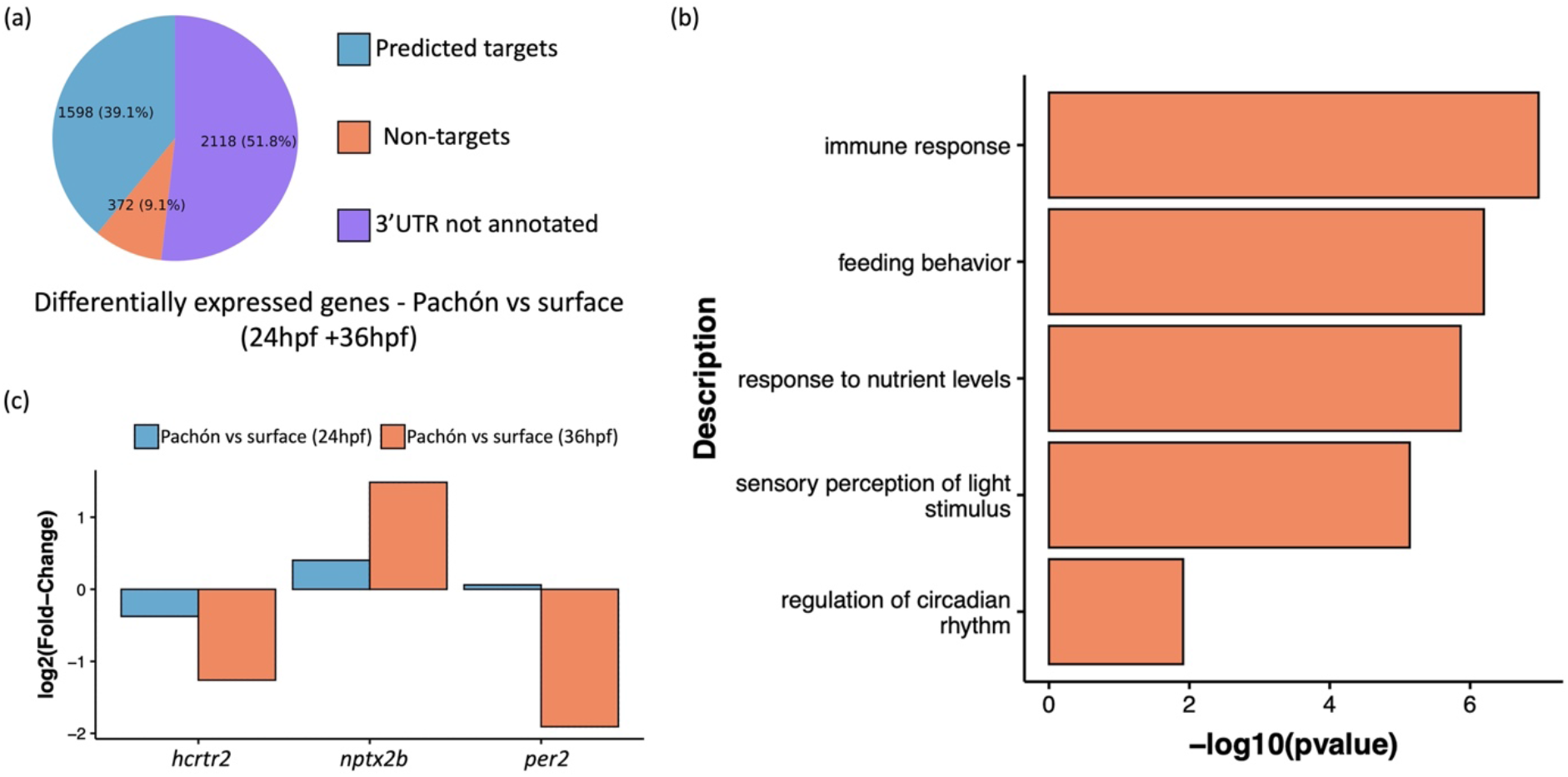
Differential expression of predicted targets of cave-specific miRNA. (a) Pie-chart depicting the percentage of differentially expressed genes which are also predicted to be targets of cave-specific miRNA, when Pachón is compared to surface fish at 24hpf and 36hpf. (b) Selected pathways from GO term analysis of all predicted and differentially expressed genes (1,598) representing biological processes associated with cave-adaptation. (c) Fold change of predicted targets, per2, hcrtr2, and nptx2b at 24hpf and 36hpf.

## Discussion

In this study, we present, for the first time, a comprehensive catalog of 683 small non-coding miRNAs for the fish *Astyanax mexicanus*. Focusing on an early stage of development, we extracted and sequenced miRNAs from 24hpf embryos of four different morphs of *A. mexicanus* – one being the extant surface fish, and the others were cave morphs of Pachón, Tinaja, and Molino. We also performed *in silico* analysis to ascertain putative 3’UTR targets of each of these miRNAs. Analyzing the small RNAseq we found that each of the cave morphs exhibit more than 100 differentially expressed miRNAs, compared to surface fish. The presence of differentially expressed miRNAs at 24hpf stage suggests that not only are the miRNA mediated silencing pathways activated at an early stage in *A. mexicanus*, but also potentially employed by cavefish morphs to facilitate cave adaptation. This idea is further strengthened by the fact that among hundreds of differentially expressed miRNAs, a small number of the miRNAs were common across all the three cave morphs, when compared to surface fish, making them essentially a cohort of cave-specific miRNAs. We investigated if these common miRNAs, through their targets, could potentially affect biological pathways associated with cave adaptation. We analyzed initially by shortlisting the best possible miRNA targets and analyzing them through GO term pipeline. Thereafter, we also cross-validated all the putative targets of the common miRNAs with a sample set of Pachón versus Surface fish mRNAseq data. Both the analysis churned out biological pathways affecting development as well as metabolism of the fish, including ones associated with oxidative metabolism, circadian rhythm, and neural development. More importantly, most of these pathways were relevant to troglomorphic adaptation, which in turn suggests a potential role of miRNA in assisting cave adaptation.

The data also indicates that numerous developmental and metabolic adaptations essential for survival in cave environments begin at the early stages of development. Especially relevant are the potential changes in the circadian and oxidative pathways so early during development.

Additionally, there is another unique observation from the dataset, which shows that the most numerous differentially expressed miRNAs were found in the Molino morph. Molino is one of the ‘younger’ morphs that colonized the caves relatively recently, approximately 110,000 years ago.^2^ Molino lacks many of the known underlying alterations associated with cave evolution that are shared by the other two cave morphs of Pachón and Tinaja.^7, 8^ This observation hints at a possible dependence on miRNAs in Molino, as a dynamic "rheostat" for fine-tuning gene networks while adapting to a relatively new and extreme environment devoid of light and nutrients. This hypothesis, while exciting, needs further testing.

Taken together, future studies into miRNAs, through gain and loss-of-functions, hold a great potential to unravel new paradigms of gene regulation, development as well as metabolism in cavefish. Our comprehensive catalogue of miRNAs in *A. mexicanus* will therefore pave the way for exploring this hitherto unexplored facet of miRNA-mediated adaptation and evolution of adaptation to new environments.

## Acknowledgments

The authors would like to thank Allison Scott and Kate Hall from the Stowers Sequencing and Discovery Genomics team for all the help with Library Preparation and Sequencing. This work is supported by Stowers Institute for Medical Research’s institutional funding, and NIH New Innovator Award 1DP2AG071466-01, NIH R24OD030214.

## Author contribution

**Tathagata Biswas:** Conceptualization; investigation; analysis; visualization; original draft preparation. **Huzaifa Hassan:** Analysis; visualization; data curation. **Nicolas Rohner:** Conceptualization; original draft preparation; funding acquisition.

## Data availability statement

Original data underlying this manuscript can be accessed from the Stowers Original Data Repository at http://www.stowers.org/research/publications/libpb-2454

## Supplementary Figures

**Figure S1:**
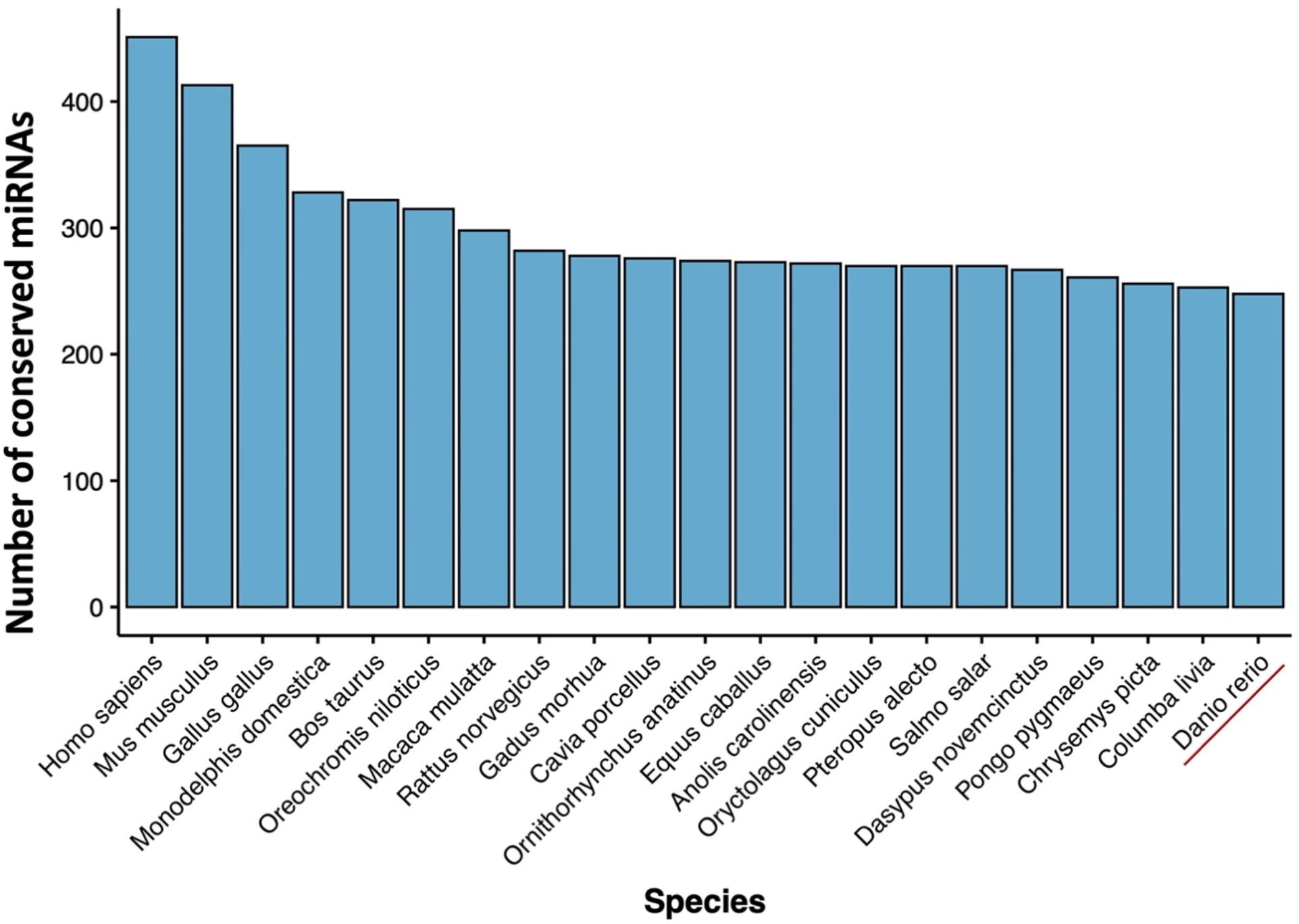
Number of A. mexicanus miRNAs conserved across miRNAs of different species (zebrafish underlined in red).

**Figure S2:**
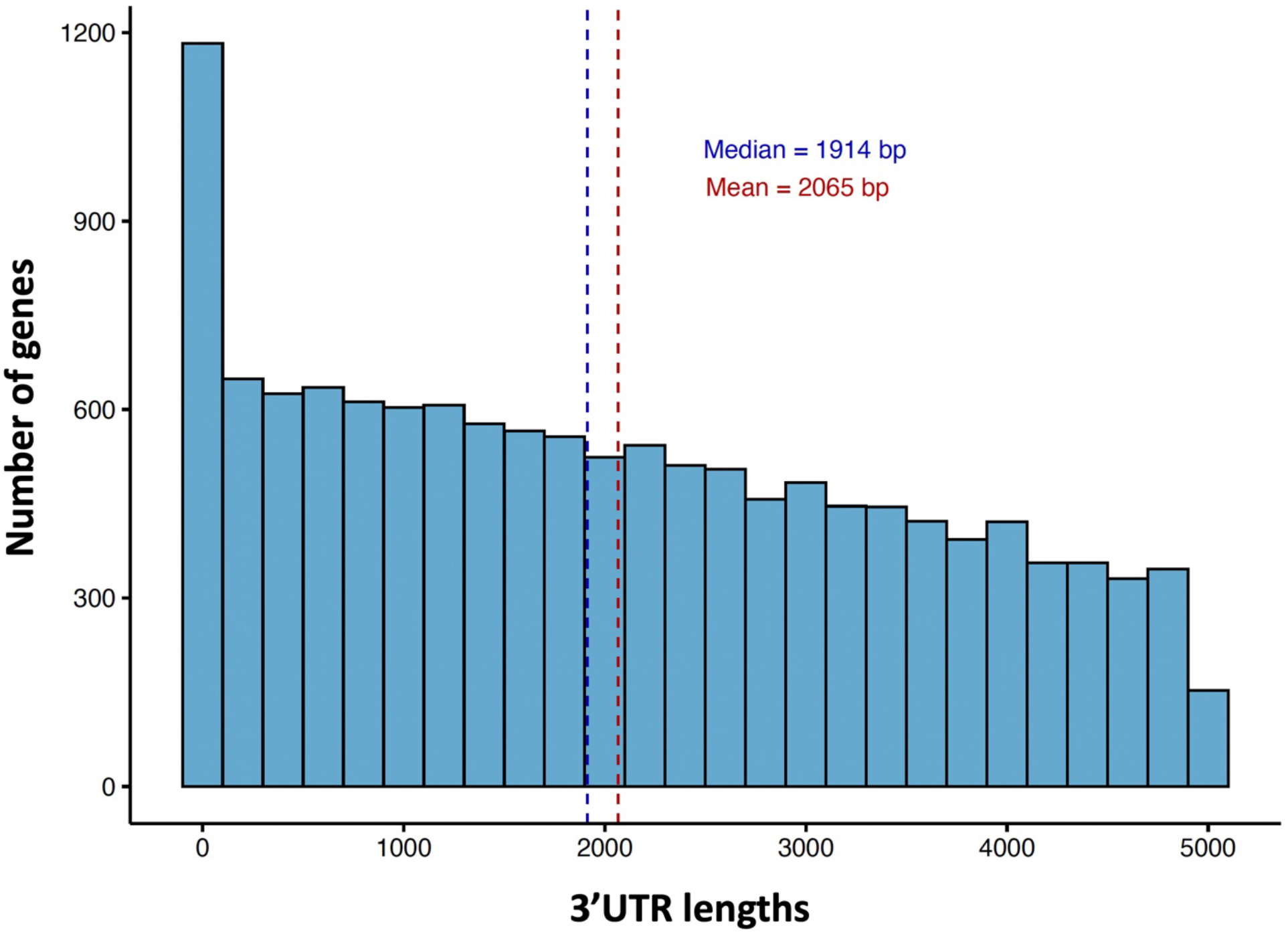
Distribution of the 3’UTR lengths of the 11,318 genes with annotated 3’UTR in Astyanax mexicanus.

**Figure S3:**
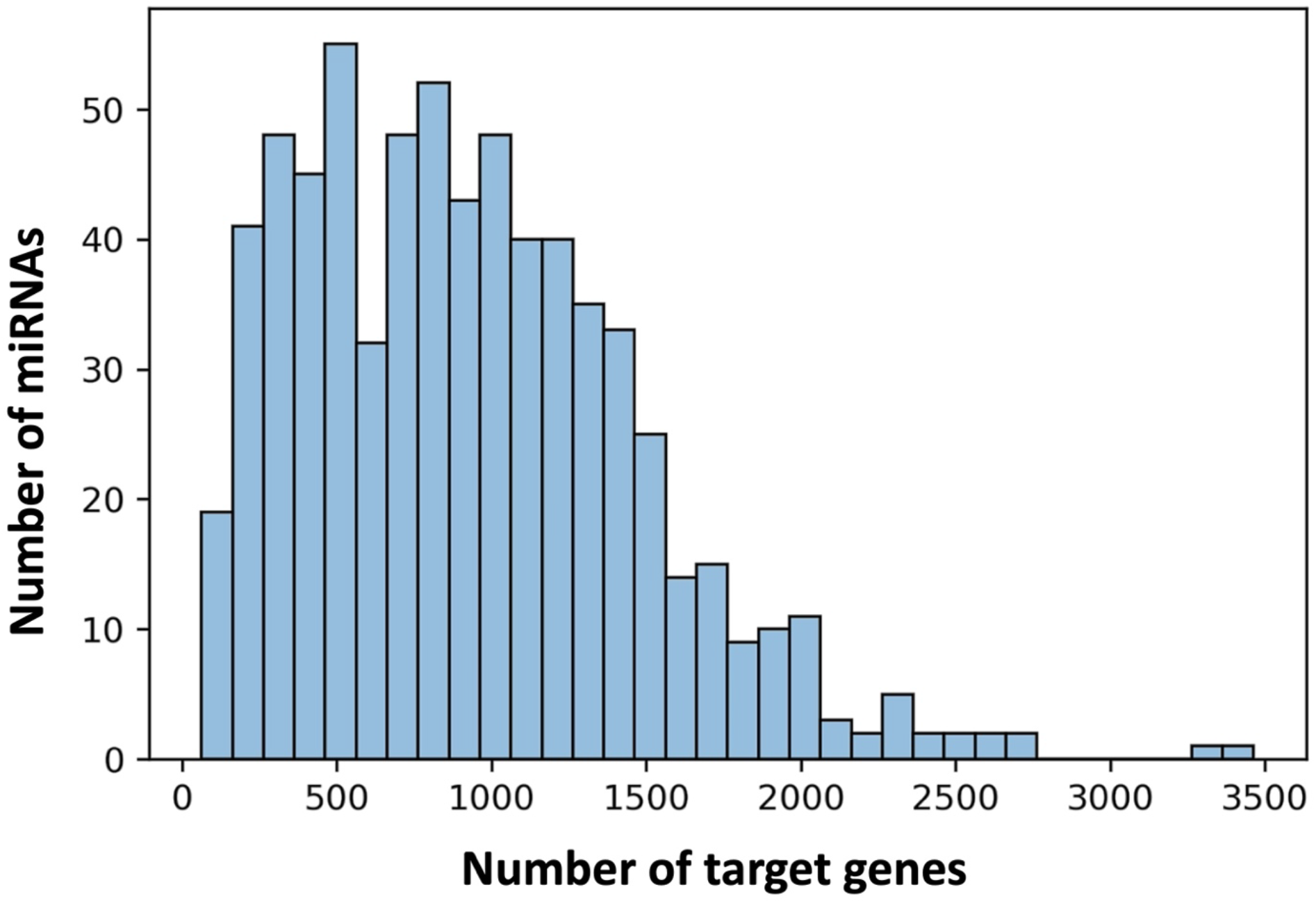
The plot depicting distribution of the number of miRNAs (Y-axis) along number of target genes (X-axis).

## References

1. Fumey J., Hinaux H., Noirot C., Thermes C., Retaux S., Casane D. Evidence for late Pleistocene origin of Astyanax mexicanus cavefish. BMC Evol Biol. 2018;18(1):43.

2. Herman A., Brandvain Y., Weagley J., Jeffery W. R., Keene A. C., Kono T. J. Y., … McGaugh S. E. The role of gene flow in rapid and repeated evolution of cave-related traits in Mexican tetra, Astyanax mexicanus. Mol Ecol. 2018;27(22):4397–4416.

3. Aspiras A. C., Rohner N., Martineau B., Borowsky R. L., Tabin C. J. Melanocortin 4 receptor mutations contribute to the adaptation of cavefish to nutrient-poor conditions. Proc Natl Acad Sci U S A. 2015;112(31):9668–9673.

4. Jaggard J. B., Stahl B. A., Lloyd E., Prober D. A., Duboue E. R., Keene A. C. Hypocretin underlies the evolution of sleep loss in the Mexican cavefish. Elife. 2018;7.

5. Olsen L., Levy M., Medley J. K., Hassan H., Miller B., Alexander R., … Rohner N. Metabolic reprogramming underlies cavefish muscular endurance despite loss of muscle mass and contractility. Proc Natl Acad Sci U S A. 2023;120(5):e2204427120.

6. Peuss R., Box A. C., Chen S., Wang Y., Tsuchiya D., Persons J. L., … Rohner N. Adaptation to low parasite abundance affects immune investment and immunopathological responses of cavefish. Nat Ecol Evol. 2020;4(10):1416–1430.

7. Riddle M. R., Aspiras A. C., Gaudenz K., Peuss R., Sung J. Y., Martineau B., … Rohner N. Insulin resistance in cavefish as an adaptation to a nutrient-limited environment. Nature. 2018;555(7698):647–651.

8. Xiong S., Krishnan J., Peuss R., Rohner N. Early adipogenesis contributes to excess fat accumulation in cave populations of Astyanax mexicanus. Dev Biol. 2018;441(2):297–304.

9. Yamamoto Y., Jeffery W. R. Central role for the lens in cave fish eye degeneration. Science. 2000;289(5479):631–633.

10. Gore A. V., Tomins K. A., Iben J., Ma L., Castranova D., Davis A. E., … Weinstein B. M. An epigenetic mechanism for cavefish eye degeneration. Nat Ecol Evol. 2018;2(7):1155–1160.

11. McGaugh S. E., Passow C. N., Jaggard J. B., Stahl B. A., Keene A. C. Unique transcriptional signatures of sleep loss across independently evolved cavefish populations. J Exp Zool B Mol Dev Evol. 2020;334(7-8):497–510.

12. Stahl B. A., Gross J. B. A Comparative Transcriptomic Analysis of Development in Two Astyanax Cavefish Populations. J Exp Zool B Mol Dev Evol. 2017;328(6):515–532.

13. Krishnan J., Seidel C. W., Zhang N., Singh N. P., VanCampen J., Peuss R., … Rohner N. Genome-wide analysis of cis-regulatory changes underlying metabolic adaptation of cavefish. Nat Genet. 2022;54(5):684–693.

14. Jonas S., Izaurralde E. Towards a molecular understanding of microRNA-mediated gene silencing. Nat Rev Genet. 2015;16(7):421–433.

15. Lau N. C., Lim L. P., Weinstein E. G., Bartel D. P. An abundant class of tiny RNAs with probable regulatory roles in Caenorhabditis elegans. Science. 2001;294(5543):858–862.

16. Lee R. C., Feinbaum R. L., Ambros V. The C. elegans heterochronic gene lin-4 encodes small RNAs with antisense complementarity to lin-14. Cell. 1993;75(5):843–854.

17. Ying S. Y., Chang D. C., Lin S. L. The microRNA (miRNA): overview of the RNA genes that modulate gene function. Mol Biotechnol. 2008;38(3):257–268.

18. Filipowicz W., Bhattacharyya S. N., Sonenberg N. Mechanisms of post-transcriptional regulation by microRNAs: are the answers in sight? Nat Rev Genet. 2008;9(2):102–114.

19. Hutvagner G., Zamore P. D. A microRNA in a multiple-turnover RNAi enzyme complex. Science. 2002;297(5589):2056–2060.

20. Lin S., Gregory R. I. MicroRNA biogenesis pathways in cancer. Nat Rev Cancer. 2015;15(6):321–333.

21. Pillai R. S. MicroRNA function: multiple mechanisms for a tiny RNA? RNA. 2005;11(12):1753–1761.

22. Cardona E., Guyomar C., Desvignes T., Montfort J., Guendouz S., Postlethwait J. H., … Bobe J. Circulating miRNA repertoire as a biomarker of metabolic and reproductive states in rainbow trout. BMC Biol. 2021;19(1):235.

23. Gebert L. F. R., MacRae I. J. Regulation of microRNA function in animals. Nat Rev Mol Cell Biol. 2019;20(1):21–37.

24. Moran Y., Agron M., Praher D., Technau U. The evolutionary origin of plant and animal microRNAs. Nat Ecol Evol. 2017;1(3):27.

25. McGaugh S. E., Gross J. B., Aken B., Blin M., Borowsky R., Chalopin D., … Warren W. C. The cavefish genome reveals candidate genes for eye loss. Nat Commun. 2014;5:5307.

26. Friedlander M. R., Mackowiak S. D., Li N., Chen W., Rajewsky N. miRDeep2 accurately identifies known and hundreds of novel microRNA genes in seven animal clades. Nucleic Acids Res. 2012;40(1):37–52.

27. Chen P. Y., Manninga H., Slanchev K., Chien M., Russo J. J., Ju J., … Tuschl T. The developmental miRNA profiles of zebrafish as determined by small RNA cloning. Genes Dev. 2005;19(11):1288–1293.

28. Desvignes T., Beam M. J., Batzel P., Sydes J., Postlethwait J. H. Expanding the annotation of zebrafish microRNAs based on small RNA sequencing. Gene. 2014;546(2):386–389.

29. Ahkin Chin Tai J. K., Freeman J. L. Zebrafish as an integrative vertebrate model to identify miRNA mechanisms regulating toxicity. Toxicol Rep. 2020;7:559–570.

30. Martin M. Cutadapt removes adapter sequences from high-throughput sequencing reads. 2011. 2011;17(1):3.

31. Chan P. P., Lin B. Y., Mak A. J., Lowe T. M. tRNAscan-SE 2.0: improved detection and functional classification of transfer RNA genes. Nucleic Acids Res. 2021;49(16):9077–9096.

32. Fu L., Niu B., Zhu Z., Wu S., Li W. CD-HIT: accelerated for clustering the next-generation sequencing data. Bioinformatics. 2012;28(23):3150–3152.

33. Robinson M. D., McCarthy D. J., Smyth G. K. edgeR: a Bioconductor package for differential expression analysis of digital gene expression data. Bioinformatics. 2010;26(1):139–140.

34. Miranda K. C., Huynh T., Tay Y., Ang Y. S., Tam W. L., Thomson A. M., … Rigoutsos I. A pattern-based method for the identification of MicroRNA binding sites and their corresponding heteroduplexes. Cell. 2006;126(6):1203–1217.

35. Dobin A., Davis C. A., Schlesinger F., Drenkow J., Zaleski C., Jha S., … Gingeras T. R. STAR: ultrafast universal RNA-seq aligner. Bioinformatics. 2013;29(1):15–21.

36. Li B., Dewey C. N. RSEM: accurate transcript quantification from RNA-Seq data with or without a reference genome. BMC Bioinformatics. 2011;12:323.

37. Wu T., Hu E., Xu S., Chen M., Guo P., Dai Z., … Yu G. clusterProfiler 4.0: A universal enrichment tool for interpreting omics data. Innovation (Camb). 2021;2(3):100141.

38. Krishnan J., Rohner N. Cavefish and the basis for eye loss. Philos Trans R Soc Lond B Biol Sci. 2017;372(1713).

39. Retaux S., Casane D. Evolution of eye development in the darkness of caves: adaptation, drift, or both? Evodevo. 2013;4(1):26.

40. Kelley J. L., Desvignes T., McGowan K. L., Perez M., Rodriguez L. A., Brown A. P., … Tobler M. microRNA expression variation as a potential molecular mechanism contributing to adaptation to hydrogen sulphide. J Evol Biol. 2021;34(6):977–988.

41. Leung A. K., Sharp P. A. MicroRNA functions in stress responses. Mol Cell. 2010;40(2):205–215.

42. Zhang J., Hadj-Moussa H., Storey K. B. Marine periwinkle stress-responsive microRNAs: A potential factor to reflect anoxia and freezing survival adaptations. Genomics. 2020;112(6):4385–4398.

43. Lunsford E. T., Paz A., Keene A. C., Liao J. C. Evolutionary convergence of a neural mechanism in the cavefish lateral line system. Elife. 2022;11.

44. Yoshizawa M., Jeffery W. R., van Netten S. M., McHenry M. J. The sensitivity of lateral line receptors and their role in the behavior of Mexican blind cavefish (Astyanax mexicanus). J Exp Biol. 2014;217(Pt 6):886–895.

45. Reiner A., Levitz J. Glutamatergic Signaling in the Central Nervous System: Ionotropic and Metabotropic Receptors in Concert. Neuron. 2018;98(6):1080–1098.

46. Traynelis S. F., Wollmuth L. P., McBain C. J., Menniti F. S., Vance K. M., Ogden K. K., . . Dingledine R. Glutamate receptor ion channels: structure, regulation, and function. Pharmacol Rev. 2010;62(3):405–496.

47. Petti A. A., Crutchfield C. A., Rabinowitz J. D., Botstein D. Survival of starving yeast is correlated with oxidative stress response and nonrespiratory mitochondrial function. Proc Natl Acad Sci U S A. 2011;108(45):E1089–1098.

48. Volpe C. M. O., Villar-Delfino P. H., Dos Anjos P. M. F., Nogueira-Machado J. A. Cellular death, reactive oxygen species (ROS) and diabetic complications. Cell Death Dis. 2018;9(2):119.

49. Pozo-Morales M., Cobham A. E., Centola C., McKinney M. C., Liu P., Perazzolo C., … Singh S. P. Starvation resistant cavefish reveal conserved mechanisms of starvation-induced hepatic lipotoxicity. bioRxiv. 2024.

50. Medley J. K., Persons J., Biswas T., Olsen L., Peuss R., Krishnan J., … Rohner N. The metabolome of Mexican cavefish shows a convergent signature highlighting sugar, antioxidant, and Ageing-Related metabolites. Elife. 2022;11.

51. Ando H., Hirose M., Kurosawa G., Impey S., Mikoshiba K. Time-lapse imaging of microRNA activity reveals the kinetics of microRNA activation in single living cells. Sci Rep. 2017;7(1):12642.

52. Hausser J., Syed A. P., Selevsek N., van Nimwegen E., Jaskiewicz L., Aebersold R., Zavolan M. Timescales and bottlenecks in miRNA-dependent gene regulation. Mol Syst Biol. 2013;9:711.

53. Beale A., Guibal C., Tamai T. K., Klotz L., Cowen S., Peyric E., … Whitmore D. Circadian rhythms in Mexican blind cavefish Astyanax mexicanus in the lab and in the field. Nat Commun. 2013;4:2769.

54. Appelbaum L., Wang G., Yokogawa T., Skariah G. M., Smith S. J., Mourrain P., Mignot E. Circadian and homeostatic regulation of structural synaptic plasticity in hypocretin neurons. Neuron. 2010;68(1):87–98.

55. Grimaldi B., Bellet M. M., Katada S., Astarita G., Hirayama J., Amin R. H., … Sassone-Corsi P. PER2 controls lipid metabolism by direct regulation of PPARgamma. Cell Metab. 2010;12(5):509–520.

56. Xiong S., Wang W., Kenzior A., Olsen L., Krishnan J., Persons J., … Rohner N. Enhanced lipogenesis through Ppargamma helps cavefish adapt to food scarcity. Curr Biol. 2022;32(10):2272–2280 e2276.

57. Biswas T., Rajendran N., Hassan H., Li H., Zhao C., Rohner N. 3D spheroid culturing of Astyanax mexicanus liver-derived cell lines recapitulates distinct transcriptomic and metabolic states of in vivo tissue environment. J Exp Zool B Mol Dev Evol. 2024.

